# Flexible neuronal participation within a reliable motor sequence

**DOI:** 10.64898/2026.07.10.737866

**Authors:** Jiahui Peng, Alison Duffy, Ilse Dippenaar, Lu Liu, Jiang Wu, Xinming Tu, Adrienne L. Fairhall, Carlos Lois

## Abstract

The reliable execution of learned motor sequences poses a challenge: they require reproducible neuron activity patterns for behavioral consistency yet must retain flexibility for modulation depending on internal states and adaptation to external contingencies. The zebra finch, a songbird, provides an ideal system to examine this problem because their vocalizations are highly stereotyped and are associated with precisely time-locked bursts in HVC, a key nucleus involved in song production. However, their songs are not immutable or totally fixed, as they can be modulated during social interactions and can recover after brain lesions. Here, we used calcium imaging of HVC projection neurons during freely produced songs to track activity across repeated renditions of the same song from seconds to weeks. We first confirmed that song-locked calcium-event timing was stable across days. Surprisingly, not every neuron showed a calcium event on every rendition of the song. We referred to the presence of such an event as participation. This participation was a stable, neuron-specific property that varied with the past and future renditions of the songs within an utterance and depended on social context. Song recovery after brain lesion and refinement during learning further revealed that neural participation could change while sequence timing remained stable. These results suggest that stable motor sequences need not rely on rigid repeated activation of the same cellular ensemble but can preserve temporal order while flexibly selecting which neurons contribute to each expression of the behavior. This provides a cellular solution to the reliability-flexibility tradeoff in learned motor control.

## Main

Learned motor sequences can be executed with remarkable precision across repeated renditions (*1*, *2*). This reliability is often linked to reproducible and temporally structured population activity in the underlying motor circuit (*3*, *4*). Yet these same circuits must also remain flexible, so that behavior can be shaped by internal and external variables, including internal motivation states, social interactions, and learning, and can be preserved or restored in response to changes caused by lesions and aging (*5*–*7*). A circuit that recruits the same neurons in the same pattern on every trial could achieve reliability but would leave little room for adaptive modulation or restoration (*8*). This tension leaves open a central question: how can a motor circuit combine reliability with the flexibility needed to adapt, learn, and recover?

Much of the current mechanistic understanding of precise motor control comes from studies of skilled movement in mammals. In trained macaques, reaching and grasping paradigms have revealed reproducible population dynamics in the motor cortex (*4*, *9*). Yet even highly practiced movements are not exact replicas at the resolution needed to compare their neural implementations. Small trial-to-trial differences in reaction time and kinematics are accompanied, and in some cases predicted, by fluctuations in firing rates of the neurons involved in the movement (*10*, *11*). Related studies in mice have further shown that uninstructed, incidental movements can account for a substantial fraction of single-trial neural variability (*12*). Such variability is informative when the goal is to identify the sources of motor error, but it becomes a confound when asking whether distinct neural activity patterns can generate essentially the same output. Resolving this ambiguity therefore requires a learned motor sequence whose repeated renditions are sufficiently stereotyped to hold behavioral output nearly constant while the underlying circuit is examined across trials and over time.

The songbird zebra finch (*Taeniopygia guttata*) offers an ideal system to study this question. Adult male zebra finches repeatedly produce renditions of the same highly reproducible song motif, defined as a stereotypical sequence of syllables (Fig. 1A). Within HVC, a premotor nucleus necessary for song production, projection neurons produce sparse bursts—brief, high-frequency clusters of action potentials—at precise syllable-locked times, and long-term imaging has shown that these preferred times can remain stable over weeks (*13*–*18*). Whether these neurons are active in every rendition, however, remains unresolved. Early single-unit recordings described song-related HVC projection neuron bursts as highly reliable across repeated motifs (*13*, *14*). However, with the electrophysiological methods used for that work, it was extremely difficult to record individual neurons beyond a few hours. Thus, the focus of that work was on investigating the timing precision of HVC neuron firing.

**Fig. 1.**
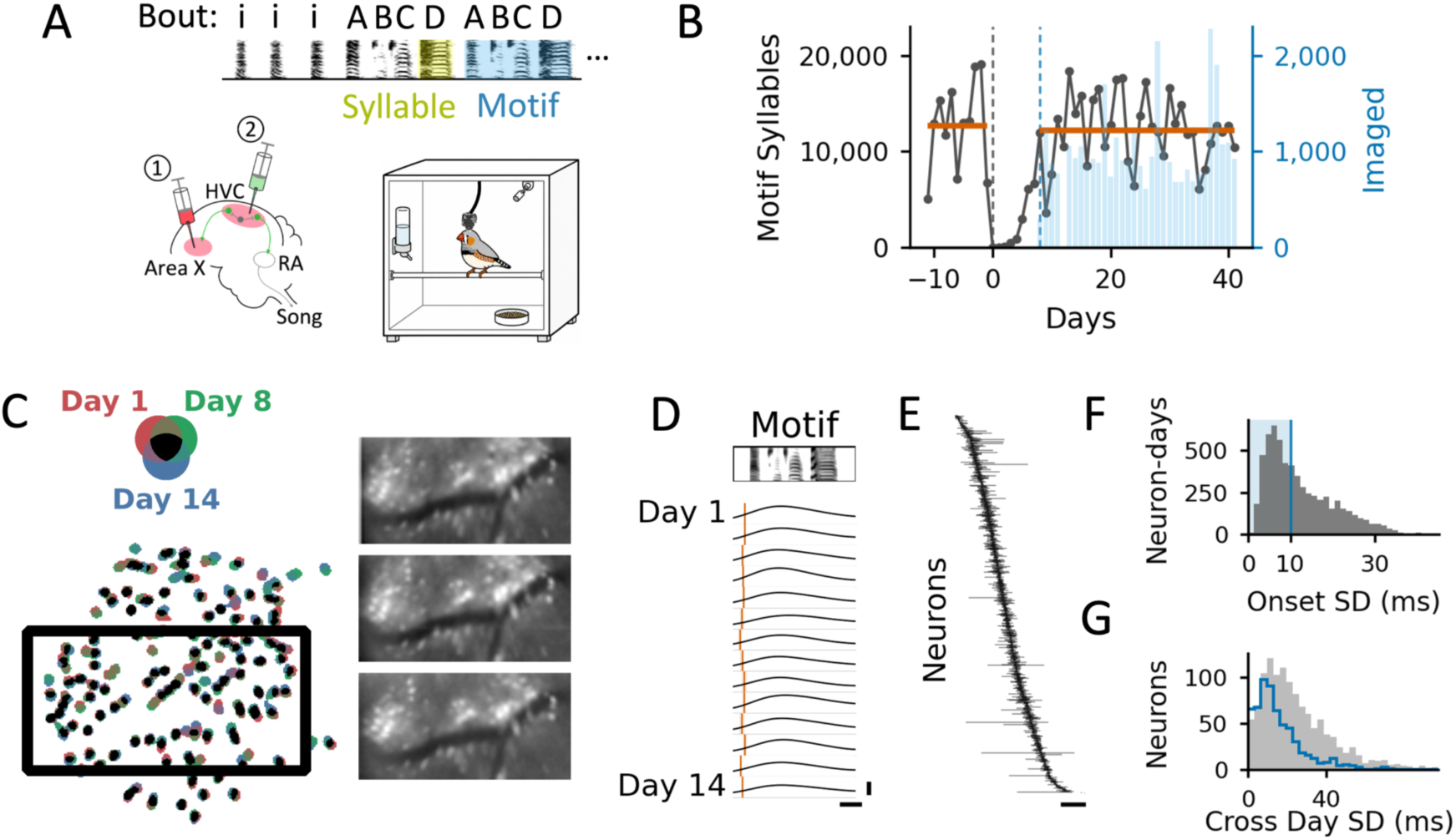
Long-term imaging and HVC event timing during song motifs. (A) Top, example of zebra finch song bout. A bout is a continuous vocalization composed of one or more introductory notes “i” in the beginning, followed by repeated song motifs containing multiple syllables, usually denoted by “ABC…”. Bottom, schematic of the recording preparation: left, ① Fluoro-Ruby injection into Area X and ② lentivirus GCaMP8m injection into HVC; right, bird implanted with a UCLA Miniscope V4 and recorded in a sound-isolation chamber. (B) Daily motif syllable counts for a representative bird. Miniscope was implanted on day 0. Gray trace, total motif syllables per day; blue bars, motif syllables recorded during simultaneous neural imaging; orange horizontal lines, mean total motif-syllable counts before implantation (12,651/day) and during the imaging period (12,217/day; 96.6% of preimplant baseline). (C) Registered fields of view and ROI masks from days 1, 8, and 14; black masks indicate ROIs detected on all three days. (D) Motif-aligned mean fluorescence traces from one example neuron across 14 recording days; orange lines mark inferred event onsets. Cross-day onset SD, 6.8 ms; onset range, 21.9 ms. (E) Sorted cross-day onset ranges for 1,204 neurons; each horizontal line spans the 25th to 75th percentile of one neuron’s onset estimates, sorted by median onset time (median IQR, 19.0 ms). (F) Distribution of day-level posterior onset uncertainty (n = 6,084 neuron-days; median +/- scaled MAD, 10.0 +/- 7.2 ms); blue line marks the 10-ms cutoff used for high-confidence timing estimates. (G) Distribution of cross-day onset SD for all neurons (gray; n = 1,204; median 20.7 ms; IQR 11.6-35.1 ms) and high-confidence neurons (blue; n = 606; median 12.6 ms; IQR 7.2–20.7 ms). Scale bars: (D) x, 100 ms; y, ΔF/F = 1; (E) x, 100 ms.

Subsequent longitudinal imaging made it possible to follow the same HVC neurons across days, but a few studies using this approach reached different conclusions. One-photon imaging in freely moving birds, including songs directed to females and solo singing, revealed substantial day-to-day changes in neuron activity (*17*). In addition, in another work imaging freely singing birds some HVC neurons did not have detectable events on some song renditions (*18*). In contrast, two-photon imaging during head-fixed, female-directed, water-rewarded singing found that syllable-locked neuronal activity was nearly invariant over weeks (*15*). The opposing conclusions of these works could be due to differences in several aspects, including different behavioral settings (freely moving versus head-fixed, directed versus undirected singing), and recording methods (one- versus two- photon imaging). Thus, whether neurons involved in the production of motor sequences need to fire for every rendition of the same task remains unresolved.

Although the zebra finch songs are highly stereotypical, there are several situations in which the behavior needs to be flexible, namely modulation of song properties depending on the social context, restoration of song after brain lesion, and refinement of the song during late stages of learning (*19*, *20*). Songs show subtle differences depending on the social context in which the male sings (*20*). For example, songs sung towards a female (female-directed songs) are faster and have lower acoustic variability than when the bird sings alone (undirected songs) (*21*–*23*). It is not known how the animal is able to produce stereotypical songs with the same neurons, while at the same time being able to modulate those songs based on the social context when required. In addition, the HVC circuit needs to be malleable, as demonstrated by the fact that a lesion of HVC leads to an initial degradation of the song, followed by almost complete recovery of the behavior (*24*–*26*). Finally, during the late stages of learning, the so-called “crystallization” period, the sequence of the song is largely established, while rendition-to-rendition variability progressively decreases toward adult levels of stereotypy (*27*, *28*).

How does the brain manage to combine high stereotypy with the ability to modify the songs under certain situations? To answer these questions, we used chronic calcium imaging with head-mounted miniscopes to follow the same HVC neurons while birds sang freely across days and weeks (Fig. 1A). Zebra finch song consists of repeated motifs, defined as a fixed sequence of syllables separated by silent gaps. A typical motif contains between 4 and 6 syllables, and the whole motif lasts around 500 to 800 ms. Birds usually produce multiple repeated motifs in succession, in a so-called bout. A bout usually starts with several introductory notes followed by 3 to 7 motifs and lasts a handful of seconds. (Fig. 1A). Birds sang routinely more than 2000 motifs per day under our recording conditions, and daily song production remained similar before and after miniscope implantation (Fig. 1B). Together, this depth of sampling provided the statistical power to determine whether neuronal recruitment was regulated across behavioral states and experimental conditions.

### Long-term imaging confirms stable timing in HVC neurons

We first determined whether we could follow the same neurons over days to test their syllable-locked firing times. Fields of view remained well aligned across imaging sessions, allowing individual neuronal regions of interest (ROIs) to be reliably registered and tracked longitudinally (Fig. 1C). We estimated neuronal burst onsets for each recording day separately using a Markov chain Monte Carlo (MCMC)-based Bayesian deconvolution algorithm adapted from (*15*). For 1,204 neurons across 7 birds that were tracked over 2-32 days, a total of 6,084 daily observations (or “neuron-days”) were made.

Within the neuron-days, estimation uncertainty of the deconvolved burst onset time closely matched previously reported values (median ± scaled MAD: 10.0 ± 7.2 ms; (*15*), 10.9 ± 6.4 ms; Fig. 1F). The onset of firing times showed high consistency across recording days (Fig. 1D). Across the population, neuron onsets tiled the motif, and each was confined to a narrow slice (Fig. 1E). Cross-day timing variability, defined as the standard deviation (SD) of estimated daily onsets for one neuron, was low across tracked neurons (1,204 neurons; median 20.7 ms; IQR 11.6-35.1 ms) and decreased further when restricted to high-confidence onset estimates (with estimation uncertainty < 10 ms, 606 neurons; median 12.6 ms; IQR 7.2-20.7 ms; Fig. 1G). Together, we confirmed that the firing times of HVC neurons were stable within the song motif across multiple days of imaging.

### Probabilistic neuronal participation is a stable cell-specific property

Despite the timing precision of HVC activity within motifs, we noticed that neurons with calcium events locked to specific syllables did not always produce them when that syllable was sung. (Fig. 2A). To quantify this observation, we restricted the analysis to 644 neurons that had stable timing (cross-day variability < 30 ms) and at least 20 motif renditions (median, 96; IQR, 59 to 133) per recording day. We defined for each neuron a firing window determined from its syllable-locked event timing and classified its response for each motif rendition as ON when a calcium event fell within that window, and OFF otherwise (Fig. 2B yellow window; see Methods). It is possible that the difference in participation we are observing here reflected differences in calcium-event amplitude, such that smaller events failed to cross the detection threshold. To address this, we quantified activity amplitude as the average ON - OFF peak contrast in raw dF/F. Across the same 644 timing-stable neurons, participation probability was unrelated to this amplitude contrast (Pearson r = 0.025, p = 0.53; Spearman rho = 0.009, p = 0.82. See Methods). Thus, the variation in participation probability across different neurons could not be explained by calcium event amplitude differences.

**Fig. 2.**
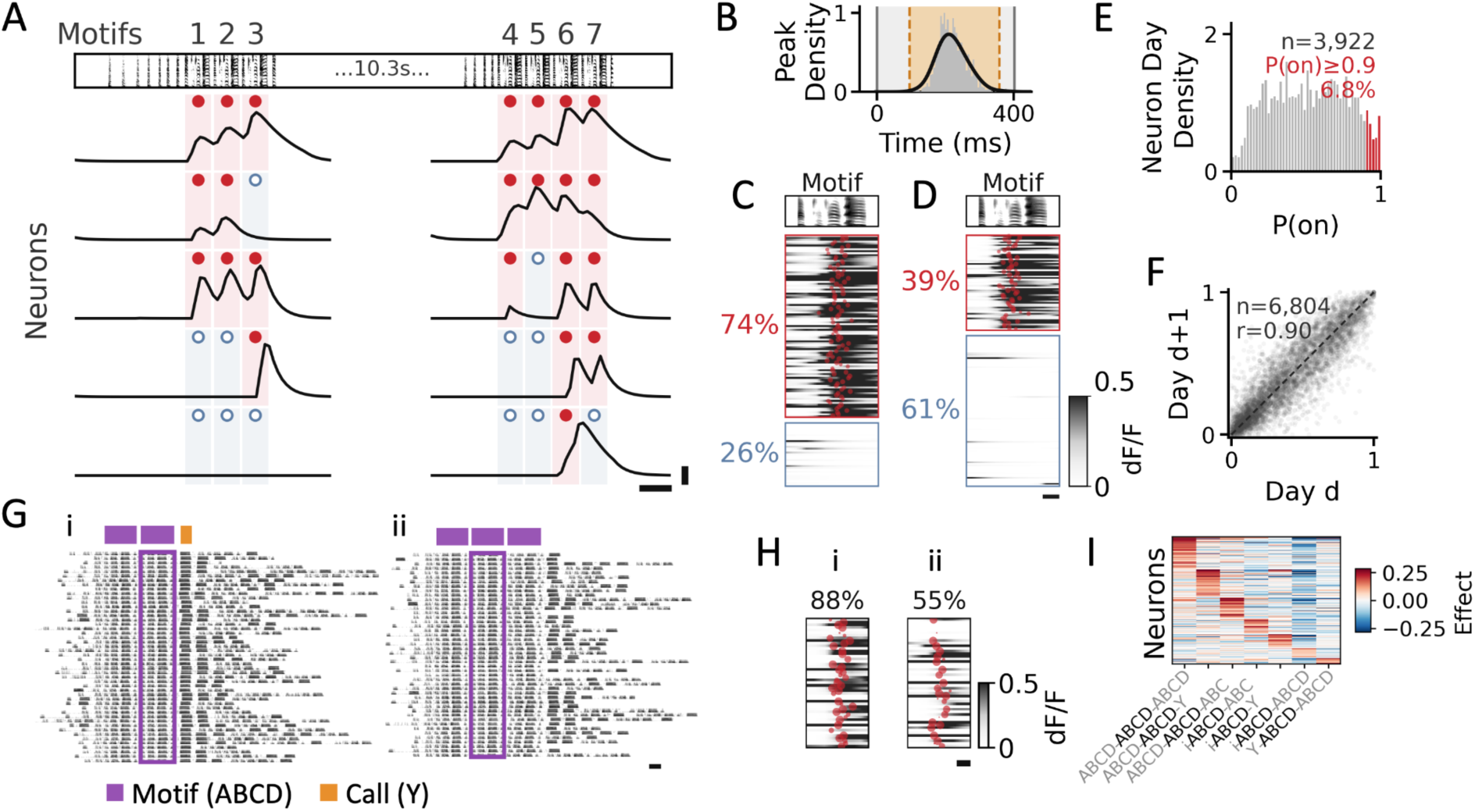
Probabilistic and syntax-dependent HVC participation in the same song motifs. (A) Example bouts with 7 total motif renditions aligned to five example neurons’ calcium traces; filled red and open blue markers indicate detected (ON) and undetected (OFF) calcium events, respectively. (B) Example participation definition for one neuron; the shaded motif window and highlighted temporal window show where calcium events were scored as ON. (C and D) Two example neurons with different participation probabilities across motif renditions. (C) was ON in 95 of 128 renditions (P(on) = 0.74), and (D) was ON in 61 of 158 renditions (P(on) = 0.39). (E) Distribution of P(on) for timing-reliable neuron-days (MCMC cross-day onset SD < 30 ms; n = 3,922 neuron-days; median, 0.52; IQR, 0.31 to 0.73). (F) Adjacent-day comparison of P(on) for the same neuron (n = 6,804 adjacent-day pairs; Spearman rho = 0.90, p < 1e-300). (G) Example syntax conditions containing the same target motif in different local sequences; i, ABCD-ABCD-Y; ii, ABCD-ABCD-ABCD. (H) Example neuron with syntax-dependent P(on) after matching trials by day and normalized bout position (i: 43 of 49 ON, P(on) = 0.88; ii: 27 of 49 ON, P(on) = 0.55; Fisher exact test, p = 6.4e-4). (I) Syntax-effect heatmap from the hierarchical participation model for one representative dataset. Rows are neurons grouped by preferred syntax, columns are syntax conditions, and colors show posterior mean syntax effects in logit units. Scale bars: (A) x, 0.5 s; y, ΔF/F = 1; (C, D, G, and H) x, 100 ms.

Among the HVC population, neurons showed a wide range of participation probabilities (Fig. 2C to E). P(on) values on adjacent days were strongly correlated (Fig. 2F), and this stability persisted in first-to-last day comparisons over longer intervals (n = 1,257 pairs separated by 3 to 32 days; median=7 days; Pearson r = 0.72, p = 1.3 x 10e-204). These results indicated that participation probability is a stable, neuron-specific property. Population-averaged participation probability was also stable: each bird’s daily mean P(on) departed from its across-day mean by a median of 0.022 (IQR, 0.014 to 0.034; Range at 2.8 to 8.9% of each bird’s mean; n = 7 birds).

The levels of neuronal participation for individual neurons were stable across multiple days. However, were there any aspects of the song that regulate the levels of participation of neurons in individual bouts? Previous work in canaries, another songbird species whose songs contain substantially more variable syllable sequences than the stereotyped motifs of zebra finches, showed that HVC activity associated with the same syllable type differed according to the sequence of syllables that preceded it, including syllables produced several steps earlier (*29*). We therefore asked whether, within the highly stereotyped zebra finch song, motif context also modulates the probability that individual HVC neurons participate in a given rendition. In zebra finch song, the same motif can occur in several distinct contexts: it may be preceded or followed by different sound elements (motif syntax; Fig. 2G), occupy different positions within a bout, and appear in bouts that vary in length and in the number of introductory notes. We therefore modeled trial-by-trial participation as a function of these song-context variables in a hierarchical Bayesian model, with recording day included as a random effect (see Methods). Participation depended significantly on motif syntax (Fig. 2G). In the example neuron shown in Fig. 2H, we compared renditions of the motif in its two syntactic contexts while matching the other context variables. The neuron participated on 88% of renditions in one syntax versus 55% in the other (Fisher’s exact test, p = 6.4 × 10e-4; Fig. 2H). After accounting for recording day, each of the song-context variables (motif syntax, bout position, introductory-note count, and bout length) contributed comparably to the remaining modulation of participation (26.5%, 28.2%, 23.9%, and 21.4%, respectively; 1,684 neurons from 7 birds). Dependence on song context was widespread across the neural population rather than restricted to a small subset of neurons. At a high-confidence threshold (probability > 0.90), motif syntax was the most prevalent individual modulator, affecting 21.7% of neurons (Fig. 2I). Other song-context variables also modulated substantial fractions of neurons, affecting 14.9-20.6% of neurons. Overall, 46.0% of neurons showed a preference for at least one context variable at this threshold, rising to 88.1% under a more permissive criterion (probability > 0.75).

We next asked whether neurons participate independently or whether they co-vary in clusters. To address this, we compared observed co-participation with a naive independent null based on each neuron’s overall participation probability. 11,430 of 42,255 pairs showed above-null co-participation (27.1%; one-sided Fisher’s exact tests, FDR-corrected q < 0.05), whereas below-null pairs were rare (16 pairs, 0.038%). We then asked whether this coordination remained after matching renditions by song context. For each neuron pair, we grouped renditions with the same recording day, motif syntax, and motif-position bin, estimated how often the two neurons would co-participate if they were independent within each matched group, and then summed this expectation across groups (see Methods). Under this context-matched null, 5,835 of 42,255 pairs showed above-null co-participation (13.8%; one-sided stratified z tests, FDR-corrected q < 0.05), whereas no pair showed significantly below-null co-participation (all FDR-corrected q > 0.254). These results indicate that rendition-to-rendition participation in HVC is not random, it is stable and cell-specific, shaped by the song context and coordinated across neurons.

### Social context reshapes HVC population states and their participation patterns

Another potential modulator of neuronal participation in HVC is social context, because zebra finch songs produce the same motifs during different social interactions and serve various communicative functions. Male zebra finches sing to females during courtship (*23*, *30*, *31*). They also sing to males, potentially associated with competition, nest guarding, or other interactions (*31*–*34*). In addition, adult males sing to juvenile males during tutoring, and social interaction with a live tutor strongly boosts vocal learning (*35*–*37*). Directed song refers to that produced toward another bird, whereas undirected song refers to that produced without an obvious recipient, usually when the bird is alone. Previous studies have shown that, compared with undirected song, female-directed singing differs in immediate-early-gene expression and neural activity in song nuclei (*38*–*40*).

To study how social context modulates HVC activity, we designed a paradigm in which each experimental session consisted of an undirected phase (15 min, bird alone in its home cage) followed by a directed phase (2 min, during which a conspecific was presented). The stimulus bird could be an adult female (female-directed, FD), an adult male (male-directed, MD), or a juvenile male (juvenile-directed, JD), presented in randomized order across sessions (Fig. 3A, top; see Methods). Head-direction analysis confirmed that under directed conditions, song production was oriented toward the presented bird (Fig. 3A, bottom). Across social conditions, the firing times of HVC neurons were syllable-locked and stable over the recording period, but they showed context-dependent differences in activity level (Fig. 3B). To test whether social context could be decoded from neural activity during individual motif renditions, we trained a leave-one-rendition-out linear decoder to classify motif renditions as undirected or female-directed, using HVC population activity (see Methods). This decoder assigns each rendition a continuous score, with higher values indicating a population state more characteristic of FD songs. Decoding was above chance in all four birds tested with held-out ROC-AUC scores ranging from 0.710 to 0.961 (Fig. 3C). This separability generalized to all pairwise context comparisons, indicating robust population state separation associated with a specific social context (Fig. 3D). We next asked whether these context-dependent shifts in population activity reflect changes in how often individual neurons participate across renditions, changes in the amplitude of their calcium signals when they do participate, or both. Participation probability was dependent on social context: one-third of neurons (306 out of 892) showed significant shifts in participation probability in at least one directed condition (Fig. 3E, left). This fraction far exceeded chance: a per-neuron label permutation null produced only ∼5% significant comparisons (median 4.7%, 95% CI 3.3 to 6.3%), and the observed fraction lay well above this null in every social context (Fig. 3E, right). In addition to changes in the levels of participation, we also observed that different social contexts also modulated the amplitude of calcium signals in individual neurons (Fig. 3F and G). We measured the relationship between participation and amplitude changes, and the correlation was very weak (Fig. 3F). In 42.4% of cases, the participation and amplitude changed in opposite directions, 25.7% with participation down and amplitude up, 16.7% with the reverse (Fig. 3G), indicating that they represent largely separable axes of modulation.

**Fig. 3.**
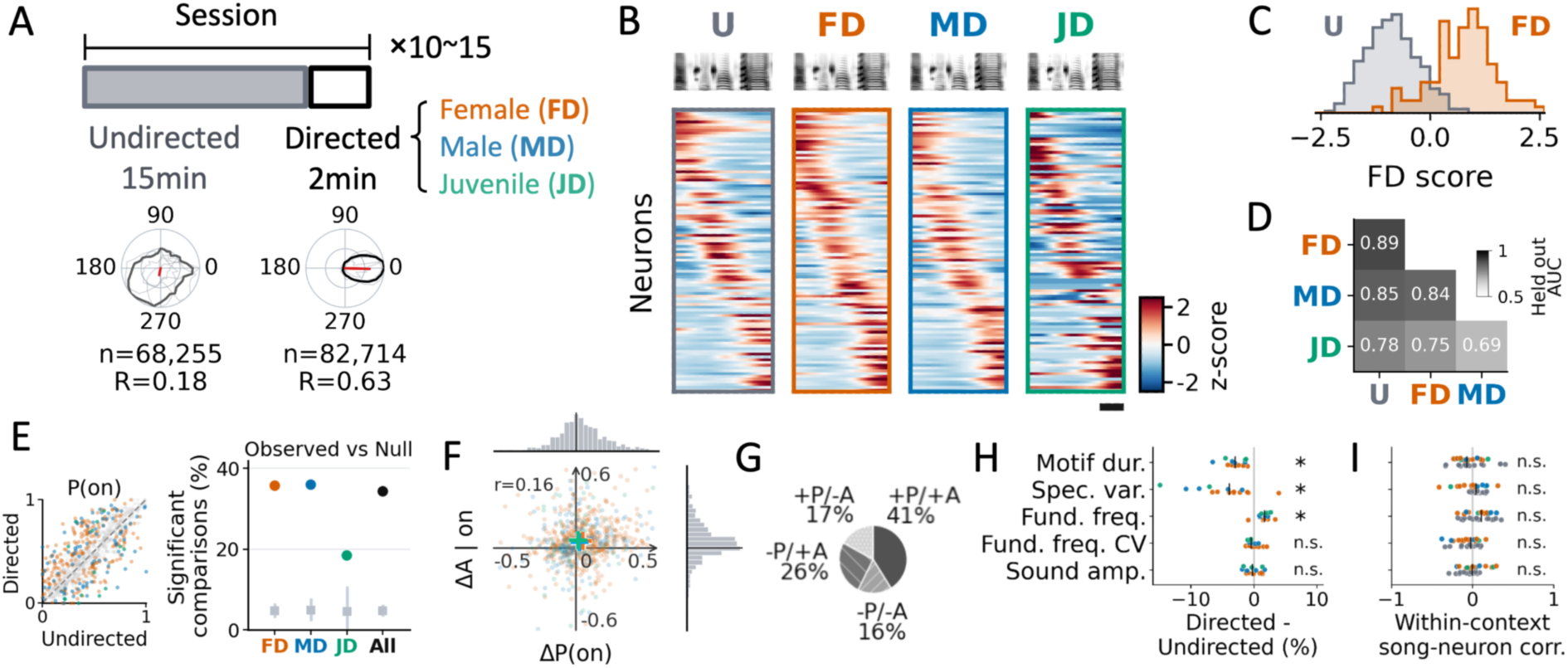
Social-context modulation of HVC activity and measured song features. (A) Social-context experimental design and head-direction analysis. Top: one session alternated undirected song (U, 15min) with directed song (2min) to a female (FD), adult male (MD), or juvenile male (JD). Bottom: polar plots show head direction during undirected and directed singing; n, number of head-orientation video frames; R, circular mean resultant length. (B) Motif-aligned population activity from an example bird across undirected and directed contexts; average motif spectrograms are shown above each heatmap, rows are 102 neurons ordered by song-locked timing, and color shows row-z-scored mean deconvolved activity. (C) Held-out decoder scores for the highest-separability FD-versus-U dataset. ROC-AUC = 0.961; 359 U and 113 FD renditions. (D) Pairwise social-context decoding matrix; entries show median held-out AUC across 9 datasets in 4 birds. (E) Left: directed versus undirected P(on) for neurons pooled across all pairwise undirected to directed context comparisons; gray points show nonsignificant comparisons and colored points show significant directed-context-modulated neurons (306 of 892 significant at p ≤ 0.05). Right: Fraction of significant neurons on the left separately for each context. Colored circles, observed fraction; gray squares with vertical lines, median and 95% CI of a per-neuron label-permutation null (1,000 iterations). Observed fractions exceed the null in every context (FD, 35.7%; MD, 36.0%; JD, 18.4%; All, 34.3% vs null median ≈ 4.5–4.8%; empirical p < 0.001 in all cases). (F) Joint distribution of social-context changes in participation probability and event amplitude on participating renditions across neuron-context comparisons. Marginal histograms show each variable separately, and plus signs mark context means across the population (FD: ΔP(on) = 0.04, ΔA|on = 0.05; MD: 0.02, 0.05; JD: 0.02, 0.12). Changes in participation probability and event amplitude were weakly correlated across comparisons (n = 773; Spearman rho = 0.16, p = 5.6e-6). (G) Quadrant summary for the joint changes in (F), grouped by the signs of Delta P(on) and Delta amplitude. (H) Directed-minus-undirected changes in measured song features; each point is one directed condition compared to undirected in one dataset (FD, orange; MD, blue; JD, green); black bars, median. n = 16 comparisons from 9 datasets in 4 birds (n = 15 for spectrogram variability). Motif duration, spectrogram variability and fundamental frequency in percent, fundamental-frequency CV in percentage points, sound amplitude in dB. Asterisks mark features with q < 0.05 from two-sided Wilcoxon signed-rank tests against zero with Benjamini–Hochberg FDR correction. Median change (q): motif duration, −3.0% (1.5e-4); spectrogram variability, −3.9% (1.9e-3); fundamental frequency, +1.7% (2.3e-4); fundamental-frequency CV, −0.41 pp (0.66); sound amplitude, −0.23 dB (0.94). (I) Within-condition, rendition-by-rendition correlations between song features and HVC decoder score; each point is one Pearson r within a single condition of one undirected-versus-directed comparison (U, dark gray; FD, orange; MD, blue; JD, green); black bars, median. Per-feature median r was tested as in (H) (n = 15 comparisons from 9 datasets in 4 birds). Median r (q): motif duration, −0.044 (0.60); spectrogram variability, 0.030 (0.66); fundamental frequency, 0.074 (0.15); fundamental-frequency CV, −0.016 (0.60); sound amplitude, 0.0003 (0.98).

Could these neural modulations be explained by song changes associated with social conditions? Previous work showed that female-directed song differs from undirected song in tempo, bout structure, and acoustic variability, with directed song generally faster and more stereotyped (*21*–*23*). Interestingly, the shifts were not restricted to female-directed songs: motif duration, spectrogram variability, and fundamental frequency changed in the same direction under male- and juvenile-directed singing as well (Fig. 3H). To test whether these song changes are associated with the HVC activity shifts across social conditions, we measured the relationship between song and neural activity on a rendition-by-rendition basis, using the same pairwise undirected versus directed comparisons as above. For each rendition, we used its decoder score (Fig. 3C, D) as a continuous readout of how directed-like the HVC population state was, together with its values for the measured song features (Fig. 3H). For each undirected versus directed comparison, correlations were computed separately within each condition, because pooling the two conditions could produce an apparent correlation driven solely by differences between contexts. The resulting correlations were small and not significant for any of the song features tested (Fig. 3I). Together, these results suggest that the context dependence of HVC participation levels is not a readout of the song features measured here but instead reflects an internal state set by the social context in which the bird is singing.

### Timing-participation dissociation persists under circuit perturbation and juvenile song crystallization

We next asked whether neuronal participation is altered in response to experimental disturbances of the brain. To this end, we lesioned the right HVC while imaging the left HVC (n=3 birds, Fig. 4A). The song degraded immediately after the lesion and then recovered over the following week (Fig. 4B). After the song recovered, calcium event peak timing remained close to pre-lesion values (Fig. 4C). Participation, however, followed heterogeneous trajectories: some neurons maintained their pre-lesion participation probability, whereas others gained or lost participation after the lesion (Fig. 4D). Participation changes exceeded matched adult drift (Fig. 4E). Thus, circuit perturbation altered cellular participation without a corresponding reorganization of the firing times, revealing a dissociation between stable HVC timing and flexible neuronal recruitment.

**Fig. 4.**
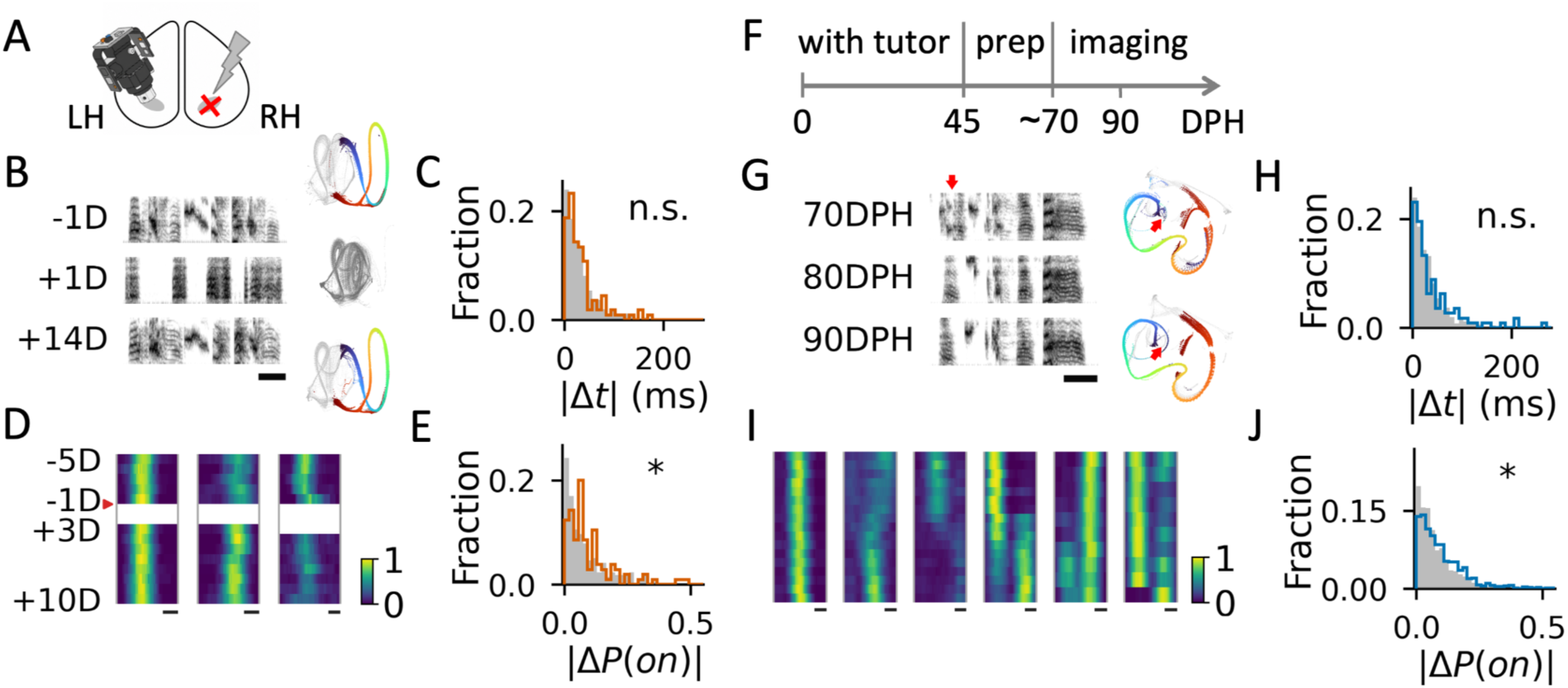
HVC timing and participation across recovery from lesion and late learning period in juveniles. (A) Contralateral lesion design, with calcium imaging in left HVC and an electrolytic lesion in right HVC. (B) Representative lesion song example from one bird. Left, spectrograms before lesion (-1 day), acutely after lesion (+1 day), and after recovery (+14 days); right, UMAP embeddings of 174-ms sliding spectrogram windows from the same days, stepped by 2.9 ms, windows that fall within the motif are colored by midpoint position (dark blue to red). (C) Distribution of absolute peak-timing shifts for individual neurons from the pre-lesion to post-lesion; the adult null was constructed from same-neuron adult pseudo-splits matched to the lesion day gap (lesion: n = 112 neurons, median, 24.25 ms; adult null: n = 523 neurons, median, 23.0 ms; one-sided Mann-Whitney U test, p = 0.459). (D) Three example neurons’ daily motif-time P(on) heatmaps; for each day and 10-ms motif-time bin, color indicates the fraction of motif renditions with at least one detected peak within +/-1 frame of that bin center (133-ms window). (E) Distribution of absolute P(on) changes between all valid pre-lesion days and post-lesion days at least 3 days after lesion, compared with center-balanced adult pseudo-splits using the same 3-day separation criterion (lesion: n = 105 neurons, median |Delta P(on)| = 0.064; adult null: n = 218 neurons, median = 0.050; one-sided Mann-Whitney U test, p = 0.020). (F) Late-juvenile experiment timeline: tutor exposure ended at 45 DPH; preparation, including surgery and miniscope weight training, continued to approximately 70 DPH; imaging began after song production returned to normal levels; 90 DPH marks the adult reference point. (G) Representative late-juvenile song example from one bird. Left, spectrograms at 70, 80, and 90 DPH; right, UMAP embeddings at 70 and 90 DPH, generated and colored as in (B). The red arrow indicates the same example syllable in the spectrograms (left) and the UMAP embeddings (right), that showed acoustic maturation over time. (H) Absolute peak-timing shifts from early to late juvenile, plotted as in (C) against the same adult null (juvenile: n = 113 neurons, median, 23.0 ms; adult null: n = 523, median, 23.0 ms; one-sided Mann-Whitney U test, p = 0.183). (I) Six example neurons’ daily motif-time P(on) heatmaps across late-juvenile days, computed and colored as in (D), illustrating stable, increased, or temporally redistributed participation. The first four neurons are from one bird recorded at 83–99 DPH, and the last two are from a second bird recorded at 78–88 DPH. (J) Consecutive-day absolute P(on) changes for late-juvenile day pairs with DPH < 90, compared with an adult-age null constructed from day pairs with 100 < DPH <= 130 using the same pooled neuron-day-pair metric (late juvenile: n = 932 neuron-day pairs; adult null: n = 1,603 neuron-day pairs; one-sided Mann-Whitney U test, p = 4.5e-11). Scale bars: (B and G) x, 50 ms; (D and I) x, 100 ms.

Does a similar timing-participation dissociation exist under natural conditions of high circuit plasticity? Late juvenile song crystallization provides such a setting. In zebra finches, the late song learning period, roughly from 60 days post-hatch (DPH) toward the adult stage (around 90 DPH), is characterized by repeated vocal practice and auditory-feedback-dependent refinement of an already recognizable song motif (*27*, *28*, *41*, *42*). During this process, syllable structure and their order in the motif are largely established, and “crystallization” refers to the gradual reduction of this variability as the song becomes adult-like and fully stereotyped (*28*, *43*). HVC premotor projection neurons are also already organized into temporally ordered song-related sequences by this stage, although these sequences continue to be refined as vocal structure matures (*44*–*46*). This allowed us to investigate whether there are changes in the stability of timing and levels of participation and whether they are decoupled as the song crystallizes.

We longitudinally imaged HVC in the same birds beginning at approximately 70 DPH, spanning the 90-DPH adult reference point, and continuing into young adulthood (Fig. 4F). Over this interval, the analyzed motifs were already recognizable, but syllable acoustic structure continued to mature (Fig. 4G). Despite this ongoing change, the firing times of HVC neurons were stable as in adult animals (Fig. 4H), indicating that the HVC temporal scaffold was already established. Participation probabilities, however, remained labile. We show six example neurons illustrating a range of trajectories, including stable participation, gradual gain, gradual loss, and changing recruitment at one or two discrete syllable-locked times (Fig. 4I). At the population level, consecutive-day participation changes before 90 DPH exceeded those observed at adult ages (Fig. 4J), consistent with progressive stabilization of participation levels during song crystallization (*44*, *46*). Together, lesion recovery and late juvenile crystallization show that the participation levels in neurons can change while preserving their specific firing times.

## Discussion

Our findings show that firing times and cellular participation are separable dimensions of the HVC sequence. Syllable-locked firing times remained stable across days, whereas their occurrence was probabilistic. Importantly, participation probability was itself a stable, neuron-specific property, systematically modulated by song and social context and coordinated across neurons. Failure of neurons to participate in a motif, therefore, did not reflect random trial-to-trial variation, but a regulated feature of the HVC population. One potential limitation of calcium imaging is that it does not directly record electrical activity. Instead, it does so indirectly by recording a side product of an action potential: an increase in intracellular concentrations of calcium. One could imagine a scenario in which neurons could have action potentials that do not translate into calcium increases. However, we believe this is a very unlikely scenario.

Song recovery after lesion and late-juvenile crystallization extend the dissociation between timing and participation from moment-to-moment modulation to longer periods, over weeks, that involve reorganization of brain circuits. These findings suggest a solution to the reliability-flexibility tradeoff in learned motor control: the stable temporal organization may serve as a scaffold within which neuronal recruitment can be flexibly reconfigured. A key open question is whether flexible recruitment contributes causally to the ability of brain circuits to learn or adapt. Addressing this question will require manipulations that selectively alter recruitment probability and measure their consequences for behavior expression, adaptation, and recovery.

## Methods

### - Animals

All procedures involving zebra finches were conducted under protocols approved by the California Institute of Technology Institutional Animal Care and Use Committee (IACUC). Birds were bred in our colony and, before experimental procedures, were housed in a large flight cage with multiple conspecifics of different ages and sexes. Following the first surgery, birds were individually housed in sound-isolation chambers under a 14 h light/10 h dark cycle for the remainder of the experiment. Juvenile birds remained in a large flight cage with breeding adults, including adult male tutors, until 45 days post-hatch, after which they were transferred to individual sound-isolation chambers.

### - Viral vector and surgery

Lentiviral vectors (LVs) were generated using standard molecular cloning procedures and produced and titrated as previously described (*47*). In all vectors, GCaMP8m (*48*) expression was driven by an internal Rous sarcoma virus (RSV) promoter. All surgical procedures for subsequent imaging were performed in the left hemisphere. Birds were anesthetized with 4% isoflurane for 1–2 min for induction and maintained under anesthesia with 1–1.5% isoflurane for the remainder of the procedure while they were head-fixed in a stereotaxic apparatus.

To retrogradely label HVC neurons projecting to Area X, a small craniotomy was made above Area X using stereotaxic coordinates referenced to the dorsal sinus (anteroposterior, 3.3-4.2 mm; mediolateral, 1.5-1.6 mm; depth, 3.5-3.8 mm). Fluoro-Ruby (10%; 100-300 nL) was injected through a glass micropipette with a tip diameter of approximately 25 μm.

Seven days after the retrograde tracer injection, a second surgery was performed to deliver the LVs and implant an imaging lens over HVC. By this time, retrogradely labeled HVC neurons exhibited strong fluorescence and could be visualized using a fluorescence stereomicroscope. Guided by stereotaxic coordinates and Fluoro-Ruby fluorescence, a 1 mm diameter craniotomy was centered over HVC. LVs were injected at 5 to 8 sites within HVC, with approximately 100 nL delivered at each site. All injections were performed at a rate of approximately 20 nL/min to minimize tissue damage. Immediately after viral injection, a durotomy was performed using a fine bent needle. A 1 mm diameter relay lens was then implanted over HVC and secured to the skull with dental cement. The exposed surface of the lens was covered with Kwik-Sil for protection.

GCaMP8m fluorescence was detectable in HVC 5 to 7 days after viral injection and lens implantation. A baseplate for a UCLA Miniscope V4 was subsequently implanted. The baseplate was temporarily attached to the Miniscope, which was positioned over the relay lens (implanted on the bird’s brain during the last surgery) while the imaging plane was optimized. Once the optimal position and orientation had been identified, the baseplate was secured to the skull with dental cement.

### - Chronic miniscope imaging

Three days after baseplate implantation, birds were fitted with a dummy scope matched to the dimensions and weight of the real Miniscope. To minimize device-related effects on behavior, the weight of the head-mounted device was counterbalanced, and human handling and other disturbances were kept to a minimum throughout the experiment. Birds were habituated to this configuration for 3–5 days. Once their song production returned to normal levels, the dummy scope was replaced with a real Miniscope.

Calcium imaging was performed using a UCLA Miniscope V4 (*49*). At the beginning of each imaging period, the focal plane was adjusted to obtain an optimal field of view (FOV), defined by the presence of a large number of neurons with sharp cellular contours. The focal position was then fixed. During consecutive recording days, the Miniscope was never removed, thereby facilitating reliable tracking of the same FOV across sessions. Images were acquired at 15–20 frames per second. Excitation light intensity was adjusted for each bird to the lowest level that provided sufficient image quality. All imaging sessions in this study were conducted within the first 4 h after lights-on, when birds typically exhibited high rates of spontaneous song production. Imaging was performed intermittently, with recording periods determined by the occurrence of spontaneous singing rather than according to a continuous recording schedule. The cumulative imaging duration was limited to 30 min per bird per day to reduce photobleaching. Data were acquired using the UCLA Miniscope acquisition software.

### - Contralateral HVC lesion

After Miniscope implantation and baseline recording of left HVC activity, adult birds received an electrolytic lesion in the contralateral right HVC. To localize the right HVC, a retrograde tracer was injected into the right Area X as described above. Seven days after tracer injection, a 3-mm-diameter craniotomy was made over the right HVC. A stainless-steel microelectrode (A-M Systems #572600; 5 MΩ impedance, 0.010 inches / 254 μm diameter) was inserted into the center of HVC to a depth of 0.5 mm. The electrolytic lesion was produced by passing a 150 μA current for 90 s. Immediately after current delivery, the lesion was visually inspected under a surgical microscope and was estimated to encompass approximately half of the fluorescently delineated HVC.

### - Song recording

Songs were recorded with high-quality microphones and digitized at a sampling rate of 44.1 kHz through an audio interface using Sound Analysis Pro 2011 software (*50*). Imaging and audio data were acquired on the same computer and temporally aligned using the timestamps associated with the two data streams.

Undirected song was operationally defined as song produced while the experimental bird was alone in the sound isolation chamber. For social context experiments, a conspecific stimulus bird, an adult female (>120 DPH), an adult male (>120 DPH), or a juvenile male (<45 DPH), was placed in a separate cage within the same sound-isolation chamber. The side of the experimental bird’s cage facing the stimulus bird was transparent, allowing visual interaction between the birds. In this study, songs produced in the presence of a conspecific stimulus bird were classified as directed songs.

### - Calcium Imaging Analysis

Image preprocessing and signal extraction: Raw Miniscope videos were converted to grayscale TIFF stacks and processed with MIN1PIPE using parameters adapted for single-photon calcium imaging in zebra finches (*51*). Videos were processed at 15 Hz without temporal downsampling and with a spatial scaling factor of 0.5. Background removal used a five-pixel structuring element, hierarchical motion correction was enabled, and neuronal seeds were selected automatically. For each detected region of interest (ROI), MIN1PIPE returned its spatial footprint and raw fluorescence trace.

Cross-day neuron registration: Spatial footprints were registered across recording days using CellReg (*52*). Fields of view were aligned using translations and rotations, and neighboring ROIs were assigned across sessions using CellReg’s probabilistic model of centroid distance and spatial-footprint correlation, with a final same-cell probability threshold of 0.5. An additional post-registration quality-control procedure was then applied. For each registered neuron, the recording session with the highest mean pairwise footprint correlation was selected as the template session. Neuron-session observations with a spatial-footprint correlation to this template below 0.7, as well as sessions in which the neuron was absent, were excluded from downstream analyses.

dF/F calculation: The fluorescence baseline, F0, was estimated separately for each neuron and recording block using a 30-s rolling median. Calcium transients were removed from the baseline estimate over two iterations. During each iteration, preliminary dF/F values exceeding the median by three noise standard deviations were masked, the mask was extended by 1 s on either side of each detected transient, masked intervals no longer than 3 s were linearly interpolated, and the rolling median was recomputed. Noise was estimated robustly from the negative half of the preliminary dF/F distribution. The resulting baseline was smoothed using a Gaussian kernel with a standard deviation of 3 s. Recording sessions at least 30 s long were processed independently. Shorter sessions were concatenated with neighboring sessions recorded on the same day when the intervening interval was less than 60 min; for an isolated short session, median fluorescence was used as a constant baseline. Corrected fluorescence was calculated as dF/F = (F − F0) / max(F0, Ffloor), where Ffloor was the 10th percentile of the distribution of median F0 values across neurons within that session.

Burst-onset estimation: Burst-onset times were estimated separately for each neuron and recording day using corrected dF/F traces from motif renditions classified as participating according to the procedure described below. The duration of each motif rendition was linearly warped to the median motif duration for that dataset. Neuron-day observations with at least 20 participating renditions were analyzed, and neurons were required to satisfy this criterion on at least two recording days. Burst onset was estimated using a Markov chain Monte Carlo (MCMC) Bayesian deconvolution procedure adapted from (*15*). For rendition j, fluorescence was modeled as yj(t) = β0j + β1j t + Aj K(t − oj; τr, τd) + εj(t), where K was a normalized double-exponential calcium kernel, oj was a latent rendition-level burst onset, and Aj was constrained to be nonnegative. Rendition-level onsets were modeled as draws from a normal distribution with a shared neuron-day onset, θ, and shared trial-to-trial temporal jitter. Rise and decay time constants and the observation-noise scale were shared across renditions within a neuron-day, whereas baseline offset, baseline slope, and event amplitude were allowed to vary across renditions. Fits were restricted to the neuron’s song-locked timing region: the peak of the modeled kernel was required to fall within the neuron’s descriptive timing slot, defined below, and the fitting interval extended 180 ms on either side of that slot. MCMC sampling was initialized using a deterministic single-kernel fit and initially performed using four chains, 1,500 burn-in sweeps, and 2,500 retained samples per chain. Fits exhibiting poor chain mixing were rerun using longer targeted sampling. Only neuron-day observations with a fitted model and an onset R-hat ≤ 1.05 were retained. The estimated burst onset for each neuron-day was the posterior median of θ, and onset uncertainty, σo, was calculated as 1.4826 times the median absolute deviation of the posterior onset samples. Cross-day timing variability was quantified as the standard deviation of the retained daily onset estimates. High-confidence analyses were additionally restricted to neuron-day observations with σo < 10 ms.

Calcium trace deconvolution and event detection: Corrected dF/F traces were deconvolved using the OASIS autoregressive model (*53*), ct = g c(t−1) + st, where ct represented denoised calcium activity and st represented the inferred nonnegative event drive. Deconvolution was performed separately for each neuron and for the same recording blocks used during baseline estimation. The decay coefficient was fixed at g = 0.70, the baseline offset was estimated under a nonnegativity constraint, the L1 penalty was used, and g was not re-estimated from individual traces. The noise scale was estimated from the negative half of each block’s corrected dF/F distribution after excluding the lowest 1% of negative values. Calcium events were defined as positive local maxima of the continuous deconvolved trace, without imposing minimum prominence or width thresholds.

Neuron participation classification: For each neuron, an initial song-locked timing center was identified from the distribution of target-motif event times using a 5-ms temporal grid and Gaussian smoothing with a standard deviation of 25 ms. When multiple events occurred during a single motif rendition, the event nearest this timing center was designated as the assigned peak. A neuron was considered callable when at least five target-motif events were detected across at least five target-motif renditions. Its descriptive timing slot was centered on the median assigned-peak time and bounded by the 10th and 90th percentiles of assigned-peak times, with a minimum width of 67 ms. A separate classification window was defined using the unexpanded 5th and 95th percentiles of assigned-peak times and was then extended by 67 ms on each side. A neuron was classified as participating in a rendition when its assigned peak fell within this classification window. Renditions containing no target-motif event, or containing an assigned peak outside the classification window, were classified as nonparticipating. Participation probability was calculated as the fraction of valid target-motif renditions classified as participating.

Amplitude-threshold controls: To test whether variation in participation reflected amplitude dependent event detection, raw dF/F traces were separated into ON and OFF renditions for each neuron. For each group, a 5%-trimmed mean trace was calculated, and peak amplitude was averaged within 33 ms of its maximum in the neuron’s timing slot. The ON - OFF amplitude contrast was then correlated with participation probability across neurons using Pearson and Spearman correlations. As a detector-independent control, ON renditions were randomly divided into two halves. One half was used to construct a baseline-subtracted raw-fluorescence template within the classification window, and the remaining ON and all OFF renditions were scored by their normalized projection onto this template. ON - OFF discriminability was quantified using ROC-AUC. At least 10 ON and 5 OFF renditions were required.

Event amplitude quantification: Event amplitude on participating renditions was defined as the height of the assigned OASIS event-drive peak. Event amplitude was left undefined for nonparticipating renditions. Kinetics-aligned raw and baseline-subtracted dF/F amplitudes were retained as parallel validation measures but were not used as the primary event-amplitude metric. In analyses of effective activity across all renditions, nonparticipating renditions were represented by an amplitude of zero so that the resulting measurement jointly reflected participation probability and event amplitude.

### - Song analysis

Song segmentation: Song recordings were segmented into candidate syllables using the pretrained multispecies WhisperSeg model (nccratliri/whisperseg-large-ms-ct2) (*54*). Song files were analyzed at a sampling rate of 44.1 kHz, with a bird-specific spectrogram time step of 2.0, 2.5, or 3.0 ms selected based on visual assessment of segmentation quality. WhisperSeg-generated syllable onset and offset times were subsequently inspected by overlaying the segmentation results on song spectrograms.

Syllable annotation: Syllable annotation was performed separately for each bird. Song waveforms were band-pass filtered between 2 and 12 kHz using a fifth-order Butterworth filter, and power spectrograms were calculated using a short-time Fourier transform with a 256-sample window and a 64-sample hop. Syllable spectrograms were converted to decibel units and represented using two complementary time-normalized matrices: one was directly resampled to 80 time bins using Gaussian-weighted averaging, whereas the other was symmetrically padded to the 75th-percentile syllable duration, or truncated when necessary, and then resampled to 80 bins. The two matrices were concatenated and flattened. The resulting vectors were reduced to 50 principal components, and a 50-nearest-neighbor graph was constructed using cosine distance. Syllables were grouped using Leiden clustering (*55*) with a resolution parameter of 1.0, and a two-dimensional UMAP embedding (*56*) was used for visualization and quality control. Cluster assignments were evaluated by examining representative syllable spectrograms, the UMAP distribution, and cluster labels overlaid on complete song spectrograms. Leiden clusters were then manually assigned bird-specific syllable identities; multiple clusters representing the same syllable type were assigned the same label, and apparent segmentation or annotation errors were manually corrected.

Song features used in the participation model: To test whether neural participation depended on song structure, four song variables were defined for each motif rendition. Motif syntax described the local song sequence. It included the song unit immediately preceding the target motif, the identity of the target motif, and the song unit immediately following it. Bout boundaries were treated as separate start and end states. Within-bout position described where the target motif occurred within the non-introductory portion of the bout. Position was normalized from 0 at the beginning of this portion to 1 near its end and divided into five equal-width bins. Time from bout onset was defined as the interval between bout onset and target-motif onset. It was categorized as 0–1, 1–2, 2–3, or ≥3 s. Introductory-note count was the number of introductory notes at the beginning of the bout. Counts were categorized as 0, 1, 2, 3, 4, or ≥5. Recording day was included as a separate categorical variable. These variables were used to model the ON/OFF participation of individual neurons across target-motif renditions.

Song features across social contexts: For Figure 3H, song features were compared between undirected song and each directed-song condition. Five song features were measured. 1) Motif duration was defined as the interval from the onset of the first motif syllable to the offset of the final motif syllable. 2) Spectrogram variability measured how much the spectrogram of a given syllable differed across renditions within the same social context. Each syllable spectrogram was restricted to 500 Hz–11 kHz and normalized to 80 time bins. Spectrograms were standardized within each dataset and syllable position. Variability was then calculated as the median pairwise distance among renditions from the same social context. A lower value indicated more consistent syllable production. 3) Fundamental frequency was measured within manually selected harmonic portions of motif syllables. These portions were defined as relative positions within each syllable so that the measurement window scaled with syllable duration. Fundamental frequency was calculated using the probabilistic YIN (pYIN) algorithm, as implemented in librosa (*57*, *58*). For each harmonic window and social context, mean fundamental frequency was calculated across renditions. 4) Fundamental-frequency CV was defined as the standard deviation divided by the mean fundamental frequency and multiplied by 100. 5) Sound amplitude was calculated separately for each motif syllable. The song waveform was band-pass filtered between 500 Hz and 10 kHz. Root-mean-square amplitude was then calculated and expressed in dBFS. Each directed condition was compared with the undirected condition from the same dataset. Changes in motif duration, spectrogram variability, and fundamental frequency were expressed as percentages relative to undirected song. Changes in fundamental-frequency CV were expressed in percentage points. Changes in sound amplitude were expressed in decibels. When multiple syllables or harmonic windows were available, their effects were summarized using the median. Thus, each point in Figure 3H represented one directed-versus-undirected comparison from one dataset.

Rendition-level song-neural correlation: For Figure 3I, each motif rendition was assigned one value for each of the five song features. 1) Motif duration was measured directly for each rendition. 2) Rendition-level spectrogram variability was calculated using a leave-one-out procedure. Each syllable spectrogram was compared with a mean template constructed from the other renditions produced in the same social context. The median template distance across motif syllables was used as the spectrogram variability value for that rendition. 3) Rendition-level fundamental frequency was defined as the median value across all valid harmonic intervals in that rendition. 4) Fundamental-frequency CV was calculated across those intervals. 5) Rendition-level sound amplitude was defined as the median amplitude across all syllables in the target motif. A pairwise social-context decoder assigned each rendition a continuous neural score based on HVC population activity. A higher score indicated that the neural population state was more similar to the corresponding directed-song condition. Pearson correlations were calculated between this neural score and each rendition-level song feature. Correlations were computed separately within undirected and directed renditions for each pairwise social context comparison. This prevented differences between social contexts from producing an artificial song-neural correlation.

### - Other quantification & statistical analysis

Hierarchical participation model: The relationship between song structure and neuronal participation was analyzed using an additive hierarchical Bayesian binomial model fitted separately for each dataset and social context. For each neuron and each observed combination of motif syntax, within-bout position, time from bout onset, introductory-note count, and recording day, the number of ON renditions, k, was modeled as k ∼ Binomial(N, p), where N was the total number of valid renditions in that combination. The log odds of participation were expressed as the sum of a neuron-specific intercept and neuron-specific categorical effects for the five variables. Intercepts were partially pooled across neurons, and each effect was scaled by a variable-specific population hyperparameter and constrained to sum to zero across its levels within each neuron. The population intercept had a Normal(0, 1.5) prior; the scale parameters for the intercept and all categorical effects had HalfNormal(1) priors, and the corresponding standardized neuron-level offsets had Normal(0, 1) priors. Neurons were required to have at least 20 valid renditions, participation probability between 0.10 and 0.99, and at least two represented levels of each song variable. Models were fitted in PyMC (*59*) using the No-U-Turn Sampler (*60*) with four chains, 1,500 tuning iterations, 2,000 retained draws per chain, and a target acceptance probability of 0.95. Convergence was evaluated using R-hat, bulk effective sample size, divergent transitions, and the Bayesian fraction of missing information. For each song variable, modulation magnitude was quantified as the posterior standard deviation of its level effects. Population-level contributions were calculated by summing posterior-mean modulation magnitudes across neurons and normalizing across the four song variables after excluding recording day. For each neuron and variable, the preferred level was the level with the largest posterior-mean effect, and evidence for modulation was quantified as the posterior probability that this preferred-level effect exceeded the mean effect of the remaining levels; neurons were classified using posterior-probability thresholds of 0.90 or 0.75 as indicated.

Pairwise co-participation analysis: Co-participation was evaluated for every unordered pair of neurons that was simultaneously callable on a common set of motif renditions. For each pair, the observed co-participation count was the number of shared renditions on which both neurons were ON. Under the naive independent null, the expected count was Np1p2, where N was the number of shared renditions and p1 and p2 were the two neurons’ participation probabilities calculated over those renditions. Positive and negative departures from this null were tested using one-sided Fisher’s exact tests applied to the 2 x 2 ON/OFF contingency table. To control for song context, shared renditions were also stratified by recording day, motif syntax, and within-bout position bin. Within each stratum s, the expected co-participation count was n1,s n2,s/ns, where ns was the number of shared renditions and n1,s and n2,s were the marginal ON counts; expected counts and their hypergeometric variances were summed across strata, and the observed-minus-expected count was converted to a z statistic. One-sided normal-tail probabilities were then calculated separately for above-null and below-null co-participation. Benjamini-Hochberg correction was applied globally across neuron pairs and separately for the two directions of each null, with q <= 0.05 considered significant. Positive calls additionally required at least three observed co-participating renditions, whereas negative calls required an expected co-participation count of at least three.

Social-context decoding: Social context was decoded separately within each imaging dataset for every available pair among undirected, female-directed, male-directed, and juvenile-directed singing. Each motif rendition was represented by a population vector containing the mean dF/F of each neuron over the motif. A binary L2-regularized logistic classifier with balanced class weights was trained using leave-one-rendition-out cross-validation. Within each cross-validation fold, neurons were included if they had finite activity measurements for at least five training renditions in each context and showed nonzero activity variation. Context pairs were analyzed only when each condition contained at least 10 renditions and at least five neurons met these criteria. The held-out decoder score was the difference between the two class logits, oriented so that larger values indicated a population state more similar to the second context in the comparison; for undirected-versus-directed comparisons, larger scores therefore indicated a more directed-like state. Pairwise separability was quantified from held-out scores using the area under the receiver operating characteristic curve (ROC-AUC), and pairwise summary matrices reported the median ROC-AUC across eligible datasets.

Social-context participation and event-amplitude comparisons: Neuronal participation and event amplitude were compared separately between undirected singing and each available directed condition within each dataset. Analysis of participation required at least 10 valid motif renditions per condition, and the participation shift was defined as ΔP(on) = P(on)_directed − P(on)_undirected. Significance was assessed independently for each neuron-context comparison by pooling the two conditions, randomly permuting their labels while preserving the original condition counts, and recomputing ΔP(on) for 1,000 permutations. The empirical probability of an effect at least as large as observed was calculated from the absolute permutation differences using a plus-one correction, and comparisons with p ≤ 0.05 were classified as context-modulated. The expected fraction of significant neurons under the null was obtained by applying the same criterion to the permuted data in each iteration. Event amplitude was analyzed only on participating renditions and was defined as the mean height of the assigned OASIS event-drive peak; a neuron-context comparison was retained when both conditions contained at least five participating renditions with finite amplitude measurements. The amplitude shift was defined as ΔA(on) = A(on)_directed − A(on)_undirected. The association between participation and amplitude shifts was evaluated across eligible neuron-context comparisons using Spearman rank correlation, and comparisons were assigned to four directional categories according to the signs of ΔP(on) and ΔA(on).

Neural-song feature correlations: To test whether measured song variation accounted for the social-context neural state, rendition-level decoder scores were related to motif duration, spectrogram variability, mean fundamental frequency, the coefficient of variation of fundamental frequency, and sound amplitude, as defined above. For each dataset and each undirected-versus-directed comparison, Pearson correlations between decoder score and each song feature were computed separately among undirected renditions and among renditions from the corresponding directed condition; at least 10 matched renditions were required for each within-condition correlation. This separation prevented the mean shift between social conditions from inducing a correlation in the absence of a within-condition relationship. For population-level inference, the undirected and directed correlation coefficients from each comparison were first collapsed to their median, and these comparison-level values were tested against zero using two-sided Wilcoxon signed-rank tests. Benjamini-Hochberg correction was applied across the five song features, with q < 0.05 considered significant.

Longitudinal timing and participation comparisons: Longitudinal timing was quantified for each neuron and recording day as the median assigned-peak time across participating target-motif renditions, with at least three assigned peaks required per neuron-day. Lesion timing shifts were calculated between the first available pre-lesion day and the last available post-lesion day, whereas juvenile timing shifts were calculated between the first day and the last day of each recording series. To construct the adult timing null, we first calculated the median interval between endpoint recordings across the pooled lesion and juvenile datasets. For each neuron in the adult control datasets, we then selected the pair of eligible recording days whose separation was closest to this interval. The absolute change in median event timing between the selected days provided one null value per adult neuron. These values were pooled to form the adult reference distribution used for both lesion and juvenile comparisons. Absolute timing shifts were compared with this null using one-sided Mann-Whitney U tests for larger experimental shifts. For lesion-related participation changes, P(on) was pooled across all valid pre-lesion days and compared with P(on) pooled across valid post-lesion days beginning at least 3 days after the lesion. A recording day was considered valid when the median target-motif trial count across neurons was at least 30; at least three valid days were required in each period, and each neuron contributed only when both pooled periods contained at least 30 renditions. The adult participation null used a center-balanced pseudo-split with the same 3-day separation and inclusion criteria. Absolute lesion-related changes in P(on) were compared with adult pseudo-split changes using a one-sided Mann-Whitney U test. For the developmental comparison, daily P(on) was calculated for neuron-days with at least 30 target-motif renditions, and absolute changes were calculated for the same neuron on consecutive calendar days. Day pairs were retained when at least 20 neurons were available. Neuron-day pairs with a midpoint age <90 days post-hatch were classified as late juvenile, whereas adult reference pairs had midpoint age >100 and ≤130 days post-hatch; the pooled neuron-day-pair distributions were compared using a one-sided Mann-Whitney U test for greater participation change in juveniles.

